# Somatomotor cortical representations are predicted by levels of short-interval intracortical inhibition

**DOI:** 10.1101/2020.04.24.057331

**Authors:** Hilmar P. Sigurdsson, Katherine Molloy, Stephen R. Jackson

**Affiliations:** School of Psychology, University of Nottingham, Nottingham UK; Institute of Mental Health, School of Medicine, University of Nottingham, UK

**Keywords:** Transcranial magnetic stimulation, somatomotor maps, short-interval intracortical inhibition, GABA

## Abstract

Transcranial magnetic stimulation (TMS) can be used to probe for the location of cortical somatomotor representations in humans. These somatomotor representations are dynamic and are perturbed following motor training, systematic intervention, and in disease. Evidence suggests that these representations are maintained by the inhibitory neurotransmitter gamma-Aminobutyric acid (GABA). In the current study, we quantified the location, outline, and variability of the first dorsal interosseous (FDI) hand muscle somatomotor representation using a novel rapid-acquisition TMS method in 14 healthy young volunteers. In addition, resting motor thresholds were measured using established protocols. TMS was also used to examine short-interval intracortical inhibition (SICI), which is thought to measure transiently activated cortical gamma-Aminobutyric acid (GABA) interneurons. Using stepwise regression, our results showed that the level of intracortical inhibition was a significant predictor of the FDI somatomotor representation suggesting that greater excitability of the hand area representation is possibly governed by greater activation of transient GABA interneurons.

---

Transcranial magnetic stimulation (TMS) can be used to construct cortical somatomotor maps which are thought to reflect the highly organised topography of the primary motor and sensory cortices [1,2]. These representations are believed to demonstrate the origin and spatial distribution of pyramidal tract neurones [2,3] representing corticospinal excitability of the stimulated muscle. Evidence shows that these somatomotor representations are dynamic, actively maintained, and likely mediated by the inhibitory neurotransmitter GABA [4]. GABA – as a mediator of neural communication – plays a pivotal role in neuroplasticity and in shaping the cortical tuning function of sensorimotor representations [5]. Furthermore, altered distribution and concentration of GABA and GABAergic mechanisms can lead to a loss of the spatial specificity of somatomotor cortical representations [6]. GABA-mediated physiological inhibition can be measured using a paired-pulse paradigm (short-interval intracortical inhibition; SICI) [7] which is thought to measure transiently activated, cortical GABA interneurons [1,2].

In this small preliminary study, we investigated how GABA-mediated cortical inhibition might contribute to the maintenance of the first dorsal interosseous (FDI) hand muscle somatomotor representation. Fourteen participants took part in our study but two had to be excluded: one participant did not show any SICI effect, and the other was excluded due to neuronavigation tracking error. Therefore, the analysis was completed with the remaining 12 participants (mean age: 17.68 ± 4.42 years, 4 females). Motor thresholds (MT) were determined individually at the start of each study session using established protocols [8]. Somatomotor maps were acquired using a rapid acquisition method [9] where MEPs are sampled randomly within a pre-defined area of the cortex. Somatomotor maps were acquired with stimulus intensity of 200% of MT up to 80 of maximum stimulator output. This study was part of a larger investigation which included acquisition of somatomotor maps of facial muscles (which exhibit a higher motor threshold compared to hand muscles) in the same study session. MEPs from the FDI muscle were time-locked and synchronised to the 3D coordinates before standardisation using a z-transformation.

The 3D coordinates were then projected to a 2D plane and re-sampled to a 35cm^2^ grid. Each pixel in the grid is appointed an approximated MEP (aMEP) based on the nearest MEP datapoint, using a triangular interpolation [10]. The variability of the FDI muscle in the map was derived from the eigenvalues (EV) of the covariance matrix, generated by fitting a 95% confidence interval ellipse. The ellipse captures the variability of the aMEPs so that when variability is low and peak aMEPs are confined to a small portion of the map the ellipse is smaller. To compare the variability between participants we computed the *area* within the ellipse using the major and minor axes of the ellipse (*Area = π x EV1 x EV2*). SICI was acquired at rest, with four subthreshold conditioning stimuli (CS) delivered to the FDI motor hotspot at 60, 65, 70, and 75% of individual MT and a test stimuli (TS) at 120% of individual MT delivered to the same location, through the same coil and an ISI of 3 ms. MEPs from each CS-TS pairing was acquired 10 times, with a total of 30 TS alone stimuli. The level of cortical inhibition was quantified with three measures: the *slope*; SICI *median* inhibition; and SICI *peak* inhibition.

The EMG signal was acquired, amplified and bandpass filtered (10-2000 Hz, with 2000 Hz sampling rate) using Brain Amp ExG and BrainVision recorder (Brain Products GmbH). TMS was delivered with two MagStim (Magstim, Dyfed, UK) stimulators connected via a BiStim module using a figure-of-eight coil coupled with a neuronavigation software (Brainsight, Rogue Research Inc., Montreal, Canada). To investigate if the area of the FDI somatomotor representation is predicted by SICI we ran a stepwise multiple regression analysis with *Area* as the outcome variable while chronological *Age, Sex, MT*, SICI *slope*, SICI *median*, and SICI *peak* were entered as predictors. Importantly, to control for the effects of any differences in chronological age, sex, and MT in our analysis, we first entered these variables in the model before entering any other variables. This analysis revealed that age (t = -2.51, b = -56.66, p = 0.04), MT (t = -2.49, b = - 23.56, p = 0.04), and SICI median value (t = -4.26, b = -1343.72, p = 0.004) were each significant predictors of the area of the FDI somatomotor representation and that the overall regression model accounted for 61% of the total variance (adj-R^2^ = 0.61, F(12,7) = 5.22, p = 0.029). Our results are presented in Figure 1.

**Figure 1.**
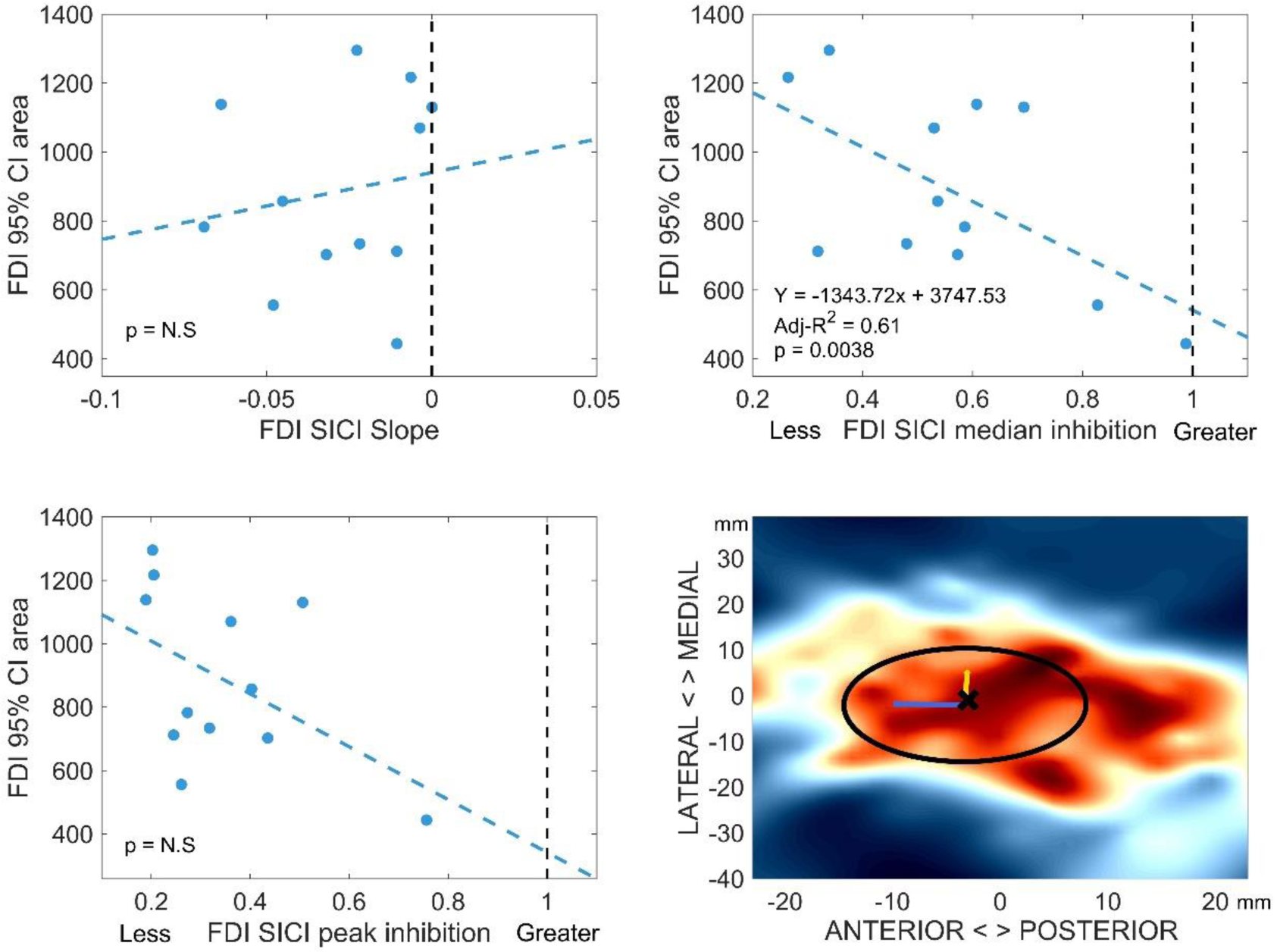
The figure demonstrates results from our study. Scatter plots are used to demonstrate relationship between SICI inhibition and FDI somatomotor map and a 2D contour plot with an example of the map from one representative participant. Dotted lines indicate the threshold between inhibition and facilitation. R^2^ indicates the total variance explained by the model. The black cross and circle in the bottom right figure indicate the centre-of-gravity and 95% confidence interval ellipse, respectively. Warmer colours represent increasing aMEP amplitude.

To summarise, our preliminary results suggest that greater excitability of the FDI somatomotor representation is related to greater activation of transient GABA interneurons.

## Acknowledgements

This work was supported by the Medical Research Council (grant number G0901321), the James Tudor Foundation, Tourettes Action (UK), and by the NIHR Nottingham Biomedical Research Centre. The views expressed are those of the authors and not necessarily those of the NHS, the NIHR or the Department of Health.

## References

[1] Siebner HR, Rothwell JC. Transcranial magnetic stimulation: New insights into representational cortical plasticity. Exp Brain Res 2003;148:1–16. https://doi.org/10.1007/s00221-002-1234-2.

[2] Chen R. Studies of human motor physiology with transcranial magnetic stimulation. Muscle Nerve Suppl 2000;23:S26–32. https://doi.org/10.1002/1097-4598(2000)999:9<::aid-mus6>3.0.co;2-i.

[3] Rothwell JC, Thompson PD, Day BL, Boyd S, Marsden CD. Stimulation of the human motor cortex through the scalp. Exp Physiol 1991;76:159–200. https://doi.org/10.1113/expphysiol.1991.sp003485.

[4] Beck S, Hallett M. Surround inhibition in human motor system. Exp Brain Res 2011;210:165–72. https://doi.org/10.1007/s00221-004-1909-y.

[5] Kolasinski J, Logan JP, Hinson EL, Manners D, Divanbeighi Zand AP, Makin TR, et al. A Mechanistic Link from GABA to Cortical Architecture and Perception. Curr Biol 2017;27:1685–91. https://doi.org/10.1016/j.cub.2017.04.055.

[6] Jacobs K, Donoghue JP. Reshaping the cortical motor map by unmasking latent intracortical connections. Science (80-) 1991;251:944–7. https://doi.org/10.1126/science.2000496.

[7] Kujirai T, Caramia MD, Rothwell JC, Day BL, Thompson PD, Ferbert A, et al. Corticocortical inhibition in human motor cortex. J Physiol 1993;471:501–19.

[8] Rossi S, Hallett M, Rossini PM, Pascual-Leone A, Avanzini G, Bestmann S, et al. Safety, ethical considerations, and application guidelines for the use of transcranial magnetic stimulation in clinical practice and research. Clin Neurophysiol 2009;120:2008–39. https://doi.org/10.1016/j.clinph.2009.08.016.

[9] van de Ruit M, Perenboom MJL, Grey MJ. TMS brain mapping in less than two minutes. Brain Stimul 2015;8:231–9. https://doi.org/10.1016/j.brs.2014.10.020.

[10] D’Errico J. Surface Fitting Using Gridfit. Matlab Cent File Exch 2005:Retrieved September 2016.

